## Supplementary Material for "Recovering *Escherichia coli* plasmids in the absence of long-read sequencing data"

### Supplementary Materials

#### Contig Alignments

For determining the composition of the bins, we used QUAST (v.5.0.2) to map the contigs of each bin to the corresponding closed reference genome. Quast uses nucmer for contig alignment. We only took contigs with a length larger than 1kb into account. Alignment of a contig was considered as correct when the sequence identity was greater than 95% and the query coverage was more than 90%, even if it was classified as “relocation” or “inversion” by QUAST (Table S4). Contigs that ambiguously aligned to several replicons were considered as correct, given that they met the previous conditions.

The alignment length of cases that were classified as ‘translocation’ by QUAST, specifying a sequence where the left and right flanking regions map to different replicons, were computed separately for all replicons.

#### Maximum theoretical recall for plasmid reconstruction

To determine the maximum recall for plasmid reconstruction from the input data (assembled short read contigs or assembly graph), we aligned either contigs or nodes extracted from the assembly graph to their respective closed reference genomes. This was important for identifying plasmid fragments that could have been missed when sequencing and which could consequently be absent in the final assembly (dead-end in the assembly graph). Therefore, these fragments could never have been reconstructed by the tools that use assembled genomes as input. It also revealed potential missmatches when mapping to the reference sequence.

Plasmids sequences were recovered with a median recall of 0.97 (IQR=0.08) when using contigs as input, and a median recall of 0.95 (IQR=0.13) was obtained when using nodes in the assembly graphs for the alignment (Figure S5 A). Notably, a total of 185 plasmids (29,3%) were perfectly recovered from contigs (recall=1), the majority of which (n=143, 77%) were small sized plasmids presenting lengths below 18 kbp. Similarly, 139 (22%) plasmids were fully recovered from assembly graphs, these were mostly small plasmids (n=122, 88%). Furthermore, 53 plasmids were fully recovered from contigs sequences only, while 7 were solely extracted from nodes in the assembly graph (Figure S5 B and C, Table S5).

Interestingly, we found that 31 (4.9%) plasmid sequences were completely missing from contigs sequences (recall=0), while 32 (5%) were missing from the assembly graph. A total of 28 plasmids (4.4%) were absent from both types of input (Figure S5 D). The sizes of these missing plasmids ranged from 763 to 4087 bp and did not contain any antibiotic resistant determinants. Interestingly, two small plasmids (Accessions: CP049974.1 and CP057228.1) were completely assembled from contigs (recall=1) but missing from the assembly graph produced by SPAdes (Table S5).

Finally, we discovered that the majority of small plasmids were contained in single nodes in the assembly graph (n=215, 79.05%) or single contigs (n=220,80.88%). Additionally, we found that 14 (5.15%) small plasmids were contained in hybrid contigs, formed by sequences derived from more than one replicon (Table S5).

### Supplementary Figures

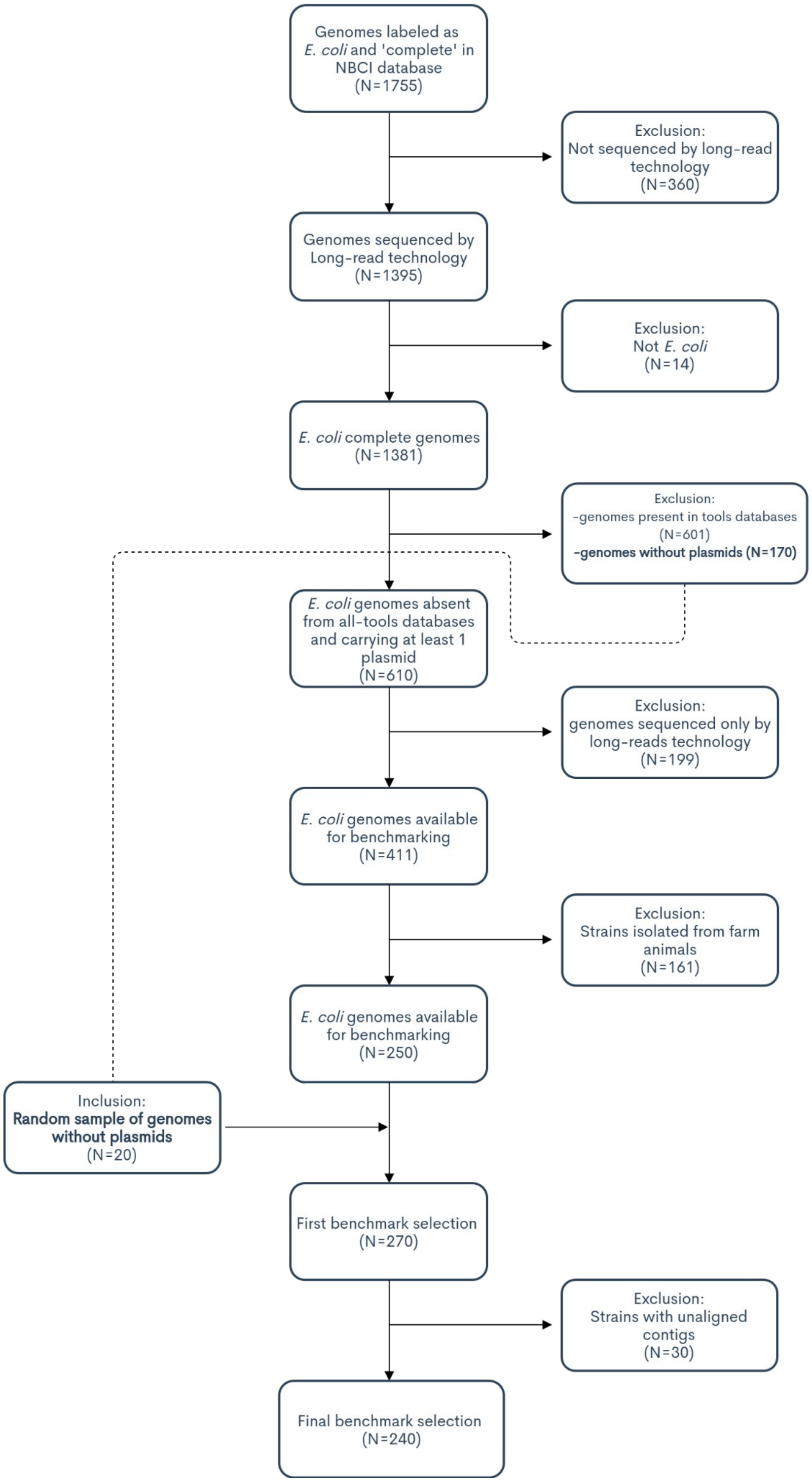


**Figure S1.** This flowchart summarizes the steps applied for selecting the 240 *E. coli* sequences included in this benchmark.

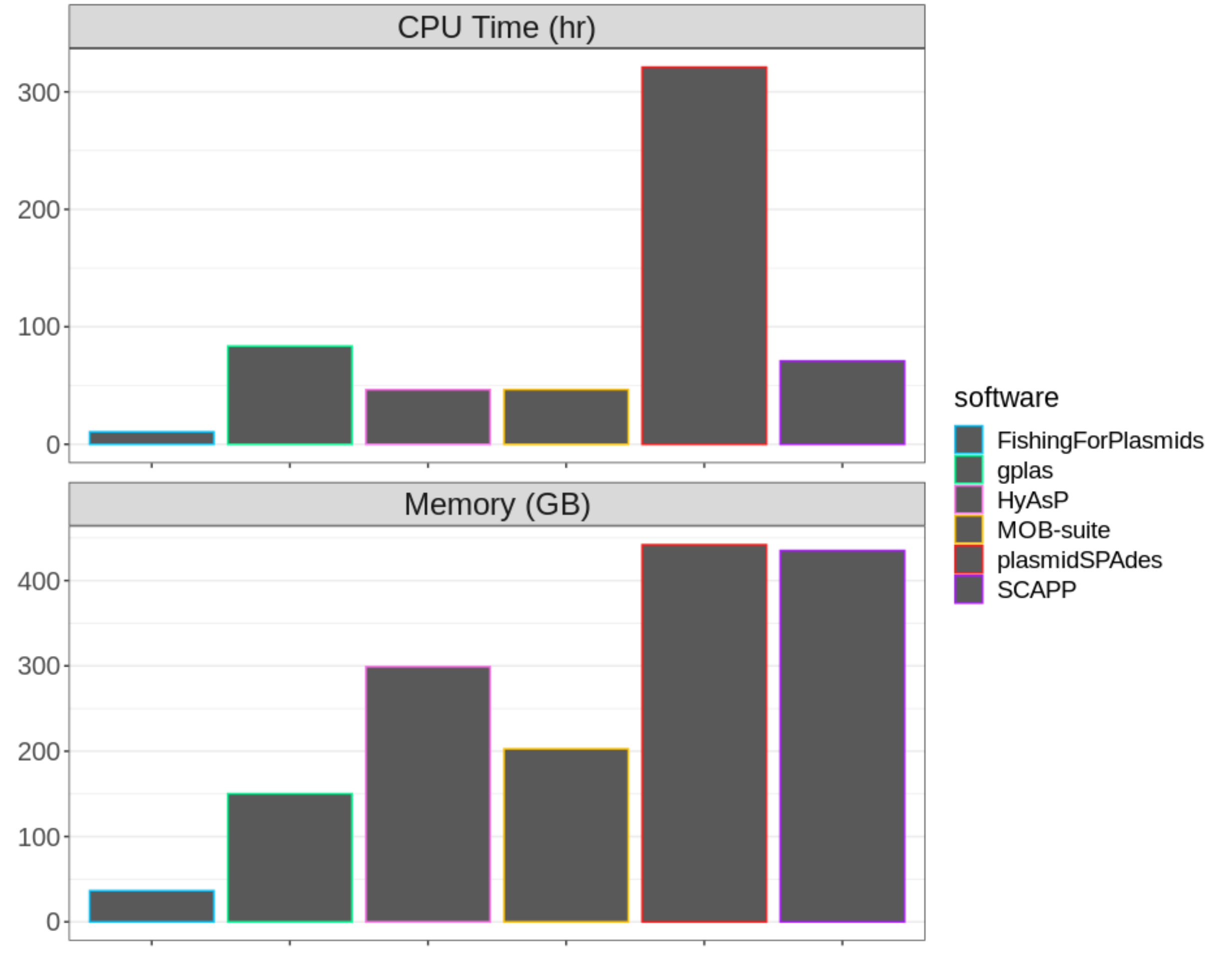


**Figure S2.** Total CPU-Time (top) and Memory (bottom) required by each tool to generate plasmid predictions in 270 *E. coli* genomes. Only 240 of these genomes were included in the final benchmark (see Methods)

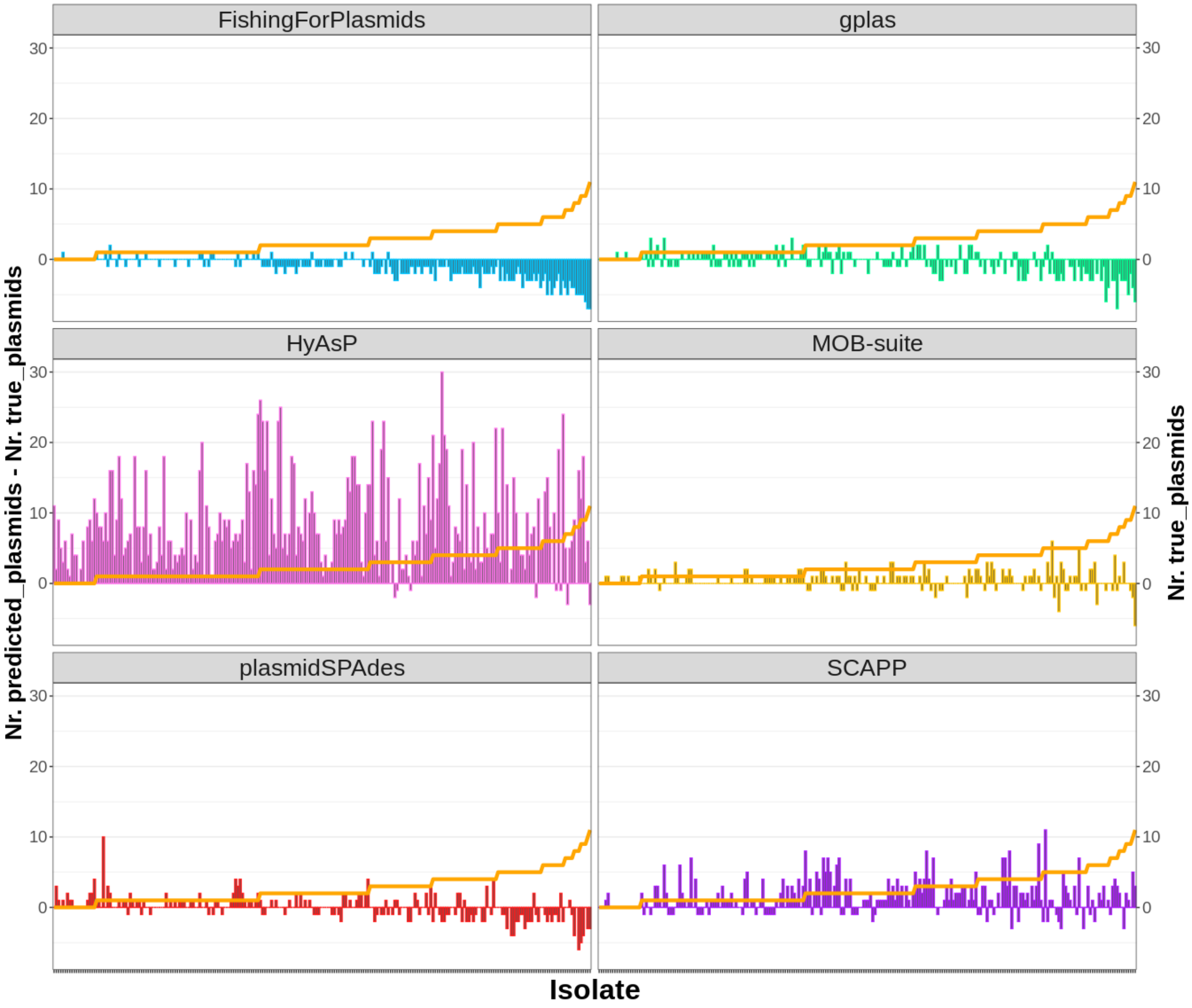


**Figure S3.** Difference between predicted and observed plasmid content per tool. Strains were ordered by increasing amount of plasmids (orange line, right y-axis). A negative value on the left y-axis indicates an underestimation of the amount of plasmids predicted for that particular strain whereas a positive value indicates an overestimation.

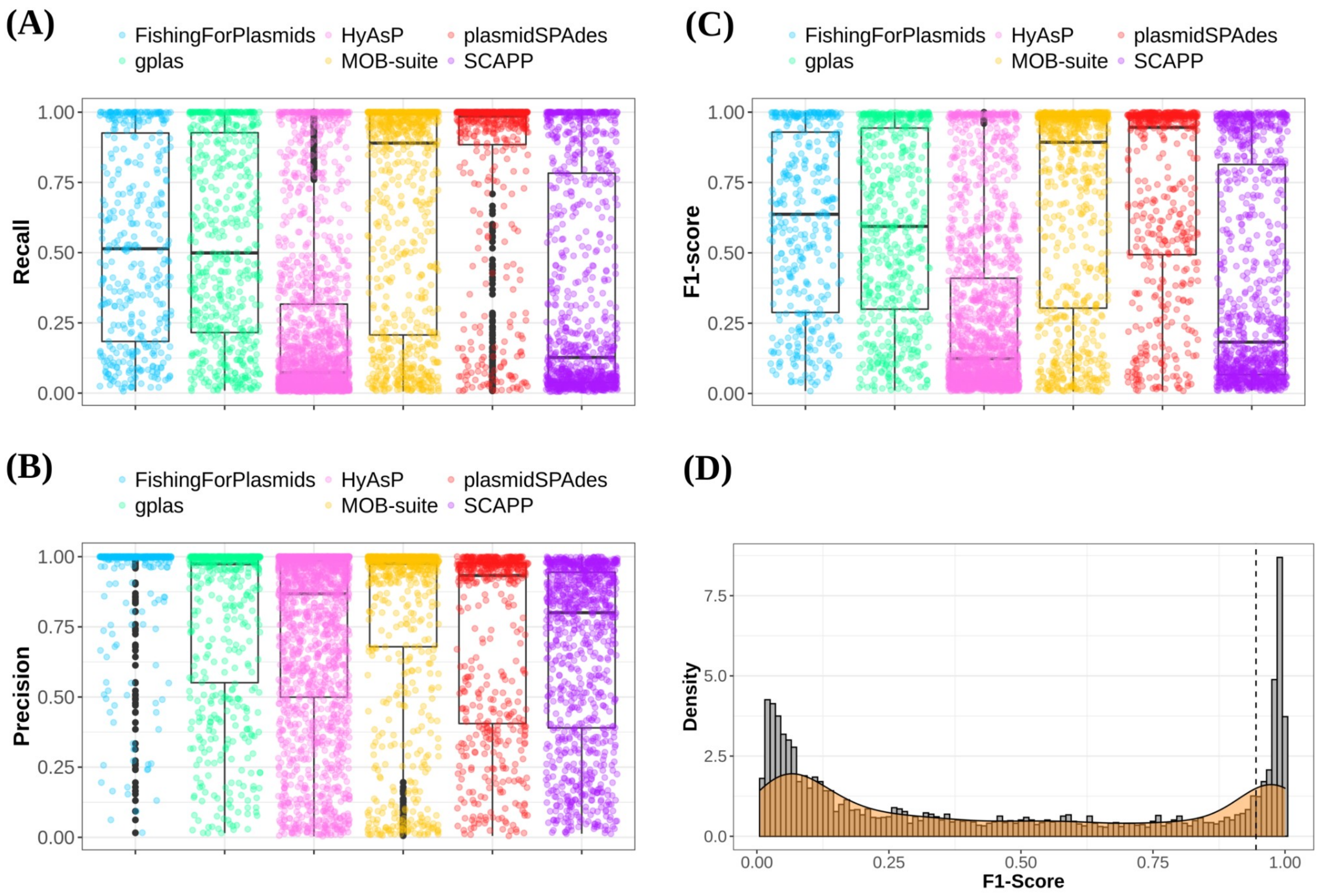


**Figure S4.** Recall (A), Precision (B) and F1-score (C) values distribution for all plasmid predictions made by each tool. (D) F1-score distribution of all plasmid predictions made by all tools combined. We established an F1-score cut-off of 0.95 (dashed line) to define a plasmid prediction as correct.

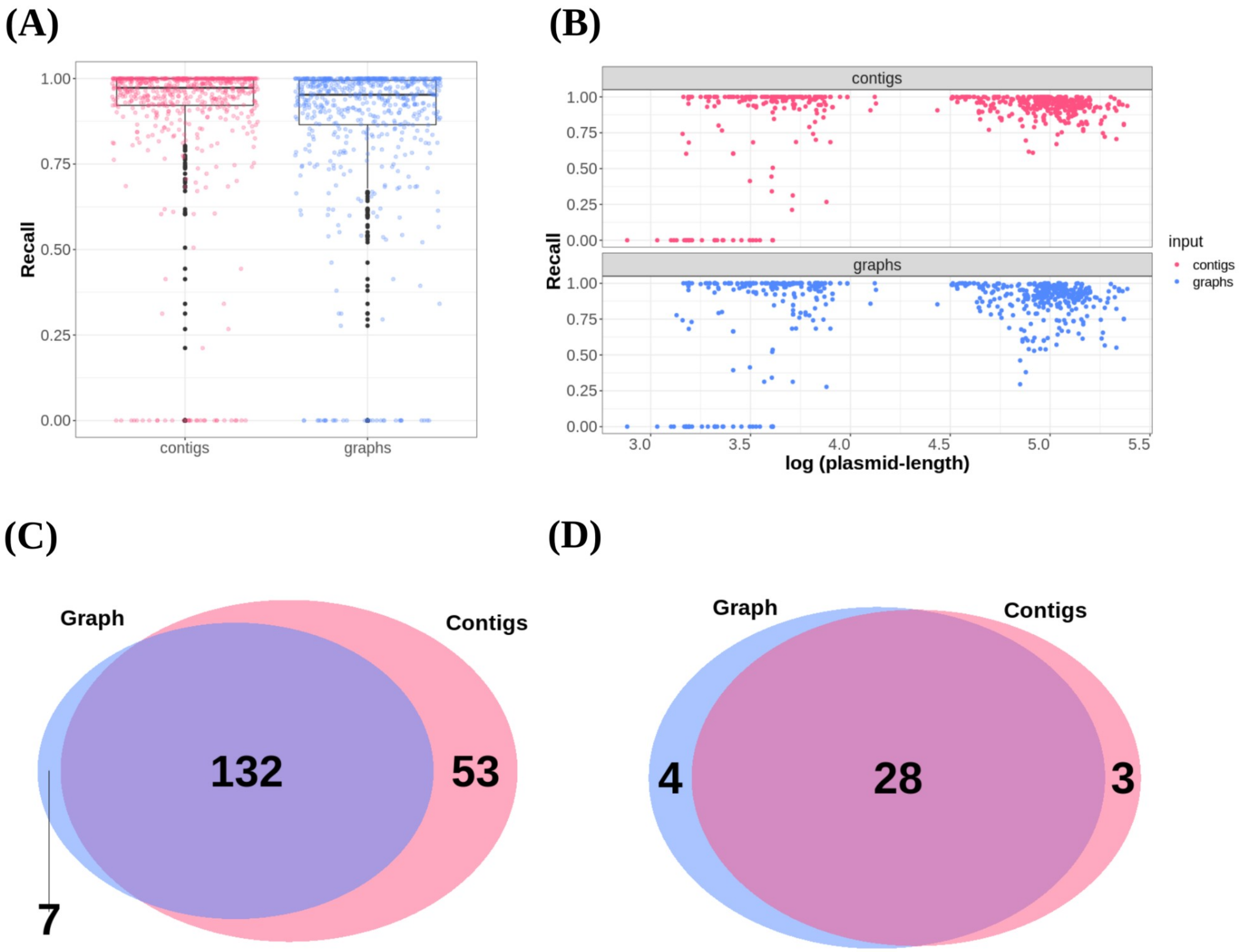


**Figure S5.** (A) Recall values obtained when aligning all assembled contigs or nodes in the assembly graph to reference plasmids. (B) Same recall values as A, as a function of plasmid size. (C) Venn diagram that shows the number of fully recovered plasmids (recall=1). (D) Venn diagram that shows completely missed plasmids (recall=0).

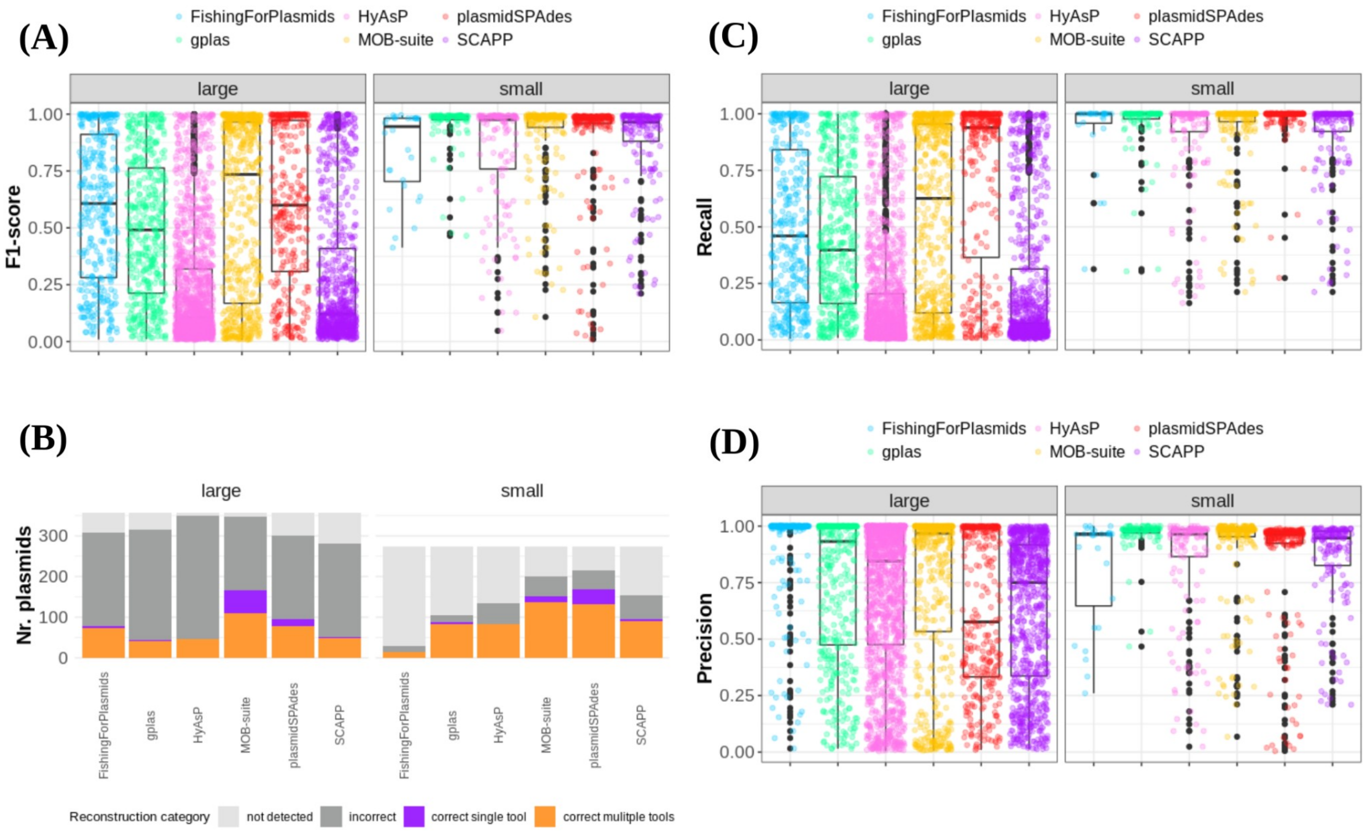


**Figure S6:** (A) F1-score value distribution for plasmid predictions according to plasmid sizes. (B) Absolute count of small and large reference plasmids that were correctly reconstructed (F1-score >0.95) - by a single or multiple tools, incorrectly reconstructed (F1-score <0.95) and not detected. (C) Recall value distribution for plasmid predictions according to plasmid sizes. (D) Precision value distribution for plasmid predictions according to plasmid sizes.

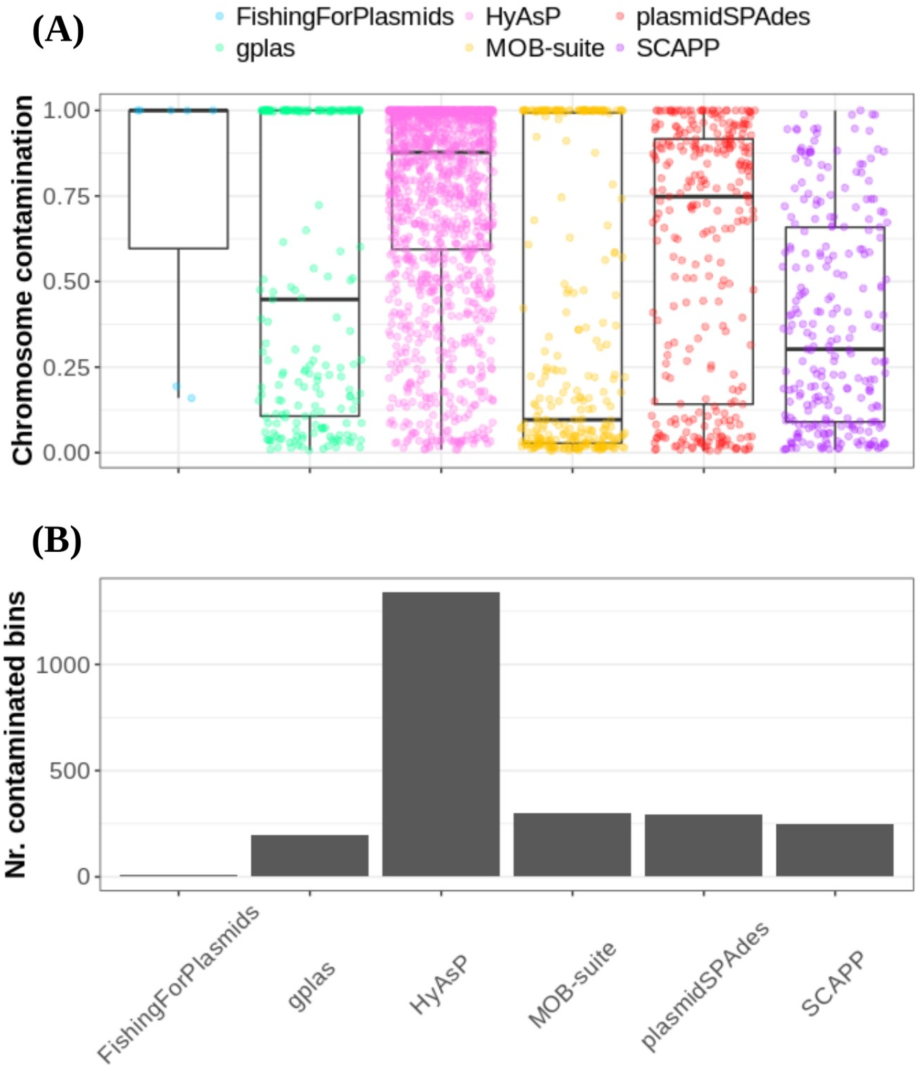


**Figure S7**. (A) Distribution of chromosome contamination values per tool. Each dot corresponds to an individual prediction that presented chromosomal sequences. (B) Count of predictions that contained chromosomal contamination.

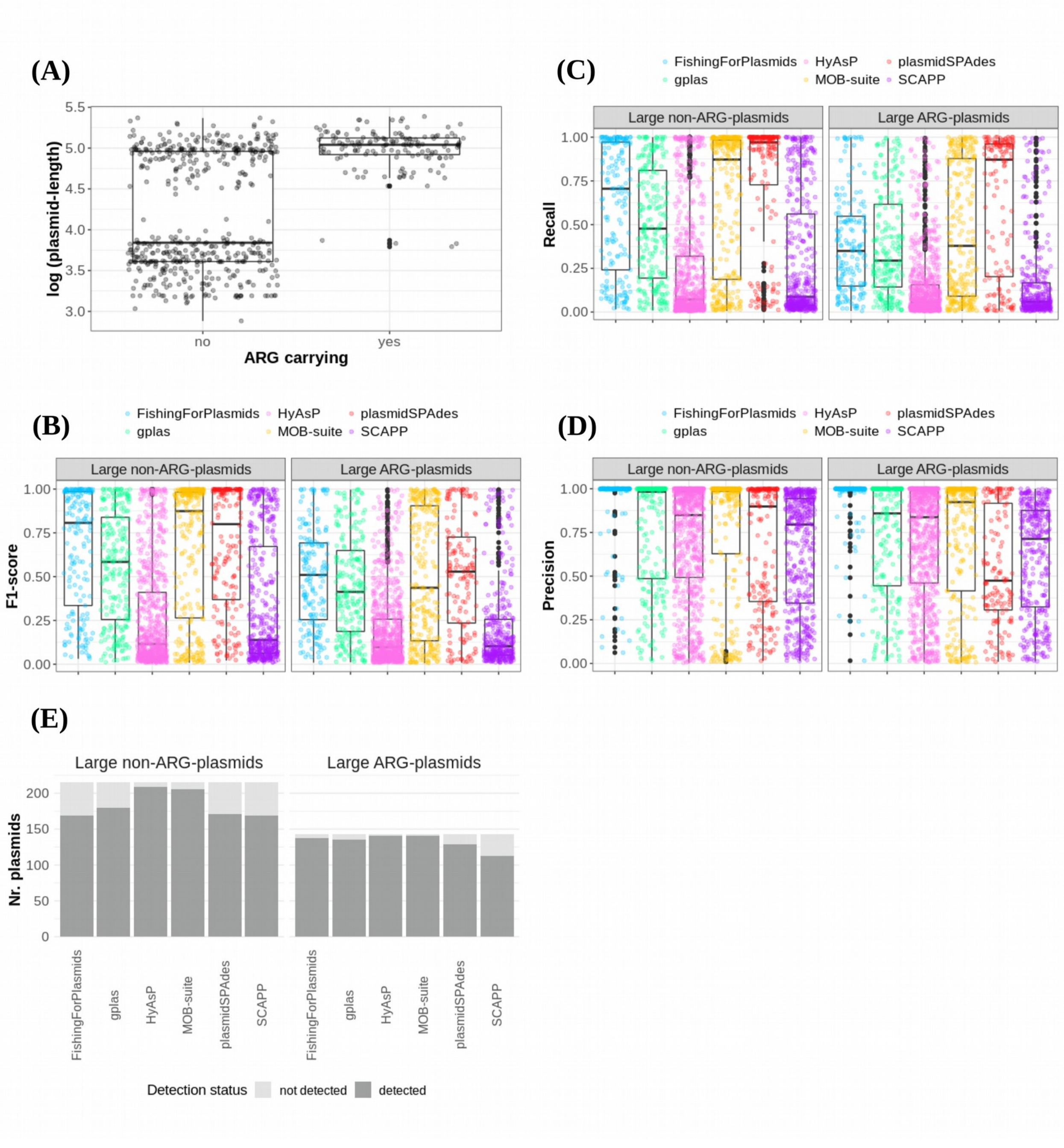


**Figure S8.** (A) Distribution of lengths for ARG and non-ARG plasmids. (B) F1-score values distribution per tool for predictions of large ARG plasmids vs. large non-ARG plasmids. (C) Recall values distribution per tool for predictions of large ARG plasmids vs. large non-ARG plasmids. (D) Precision values distribution per tool for predictions of large ARG plasmids vs. large non-ARG plasmids. (E) Bar plots showing absolute counts of detected and not detected reference plasmids.

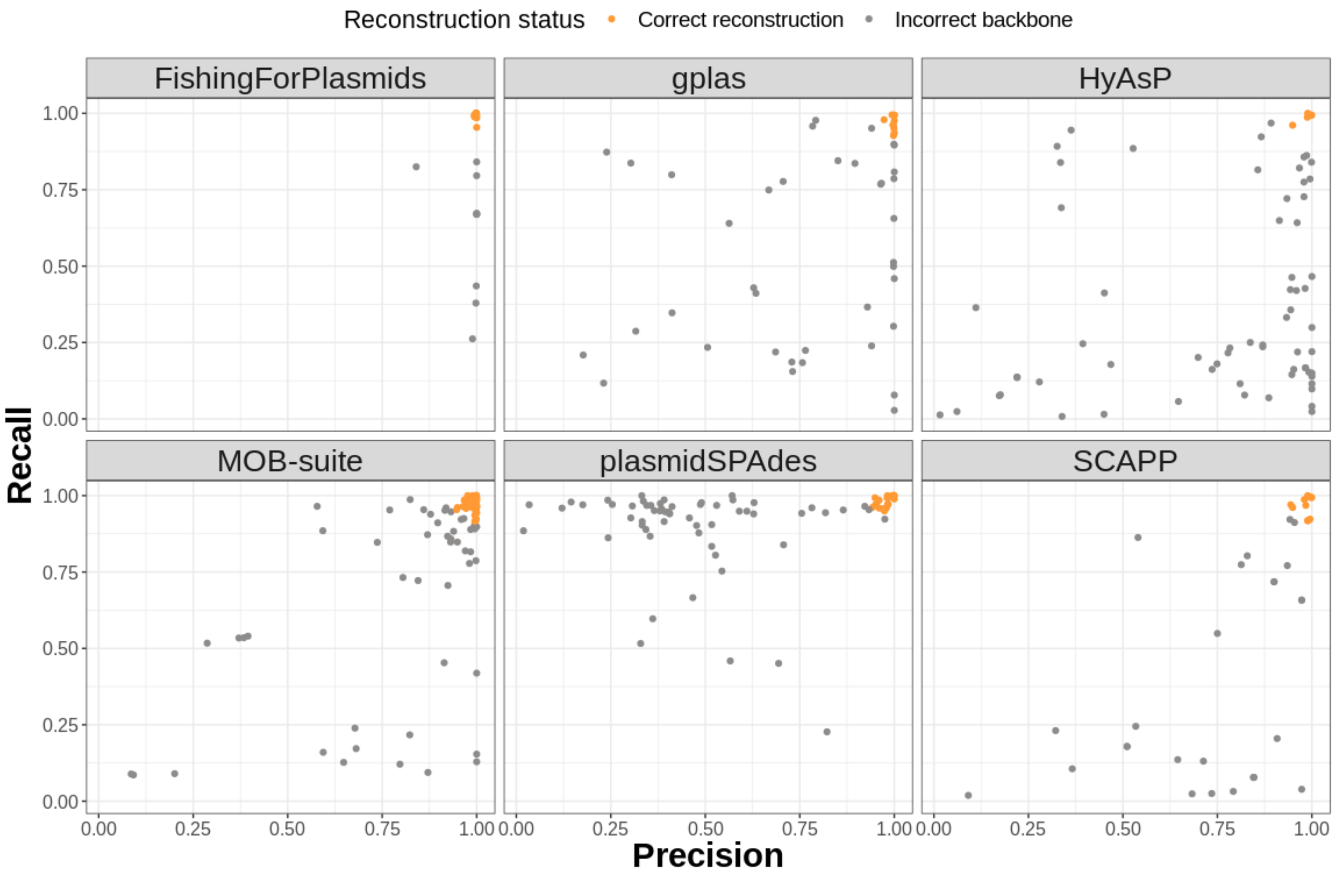


**Figure S9:** Scatter plot that shows precision (bp) and recall (bp) values for predictions that presented a Recall(ARG) equal to 1.

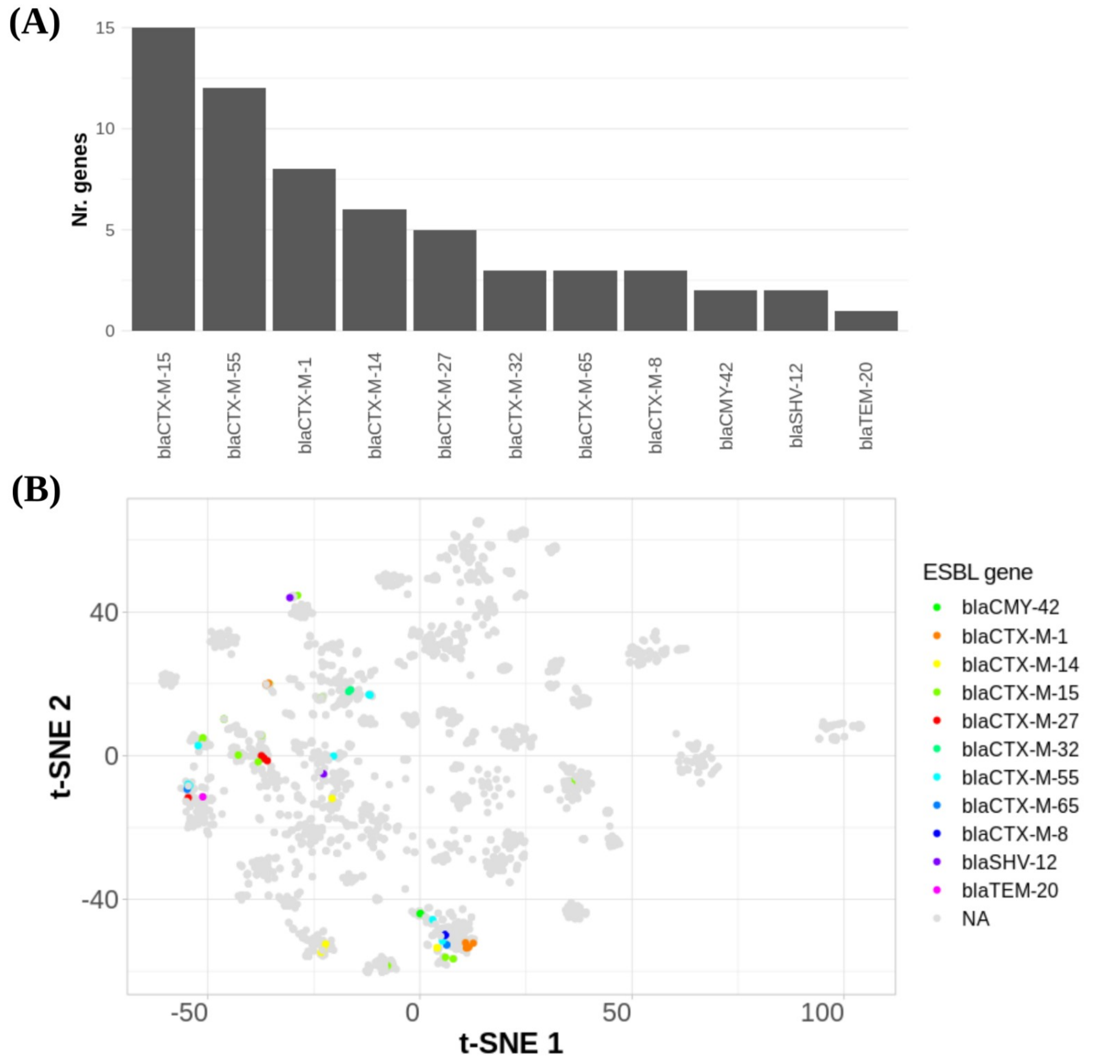


**Figure S10**. (A) Absolute count of ESBL variants in the benchmark data set. (B) tSNE created based on plasmids k-mer distances obtained with Mash (k=21, s1000). ESBL-plasmids included in the benchmark are colored according to distinct ESBL genes.

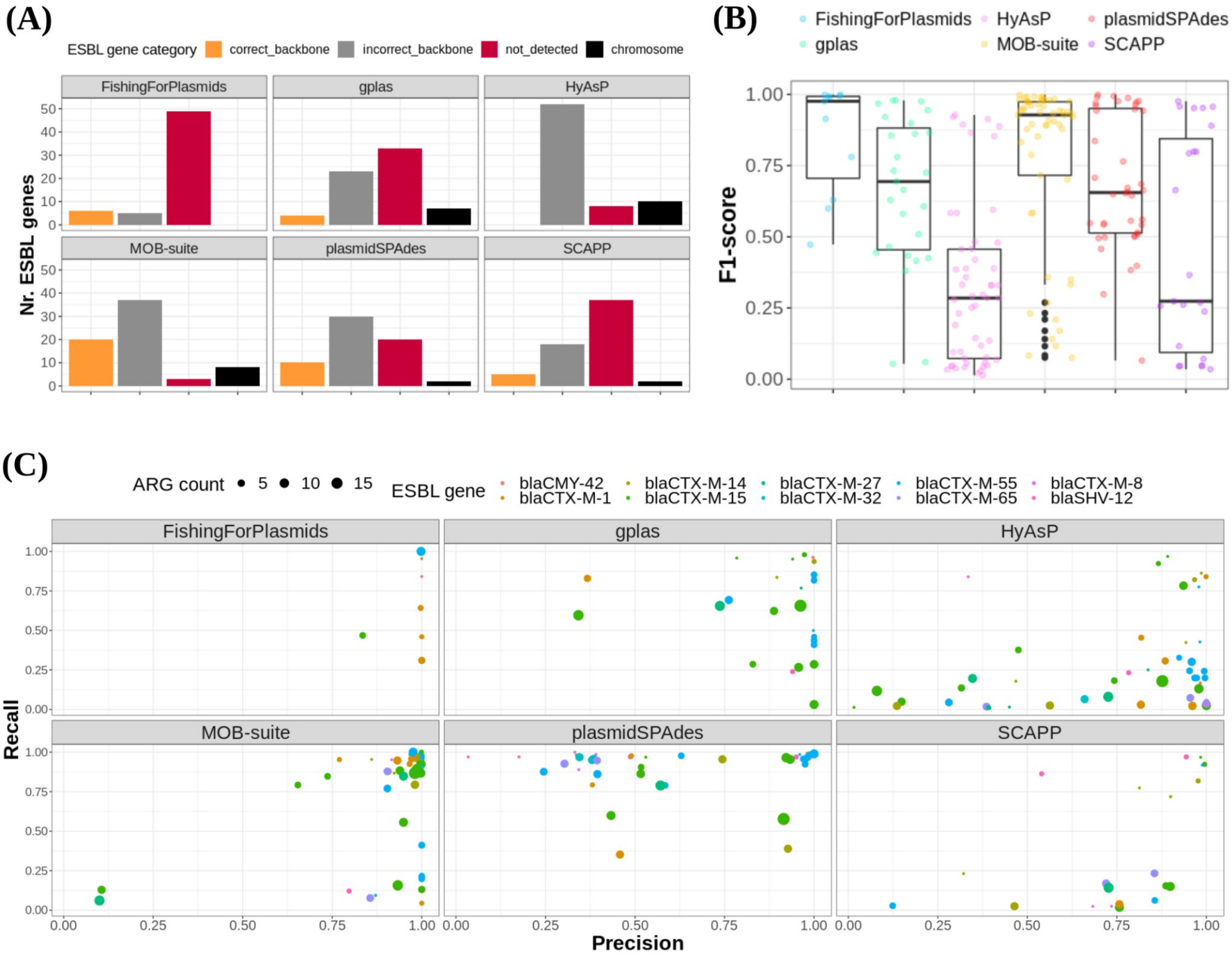


**Figure S11**. (A) Absolute count of ESBL genes according to prediction status. The different prediction status were determined according to the following criteria. Correct backbone: ESBL gene was included in a bin that presented an F1-Score >=0.95, incorrect backbone: ESBL gene was included in a bin that presented an F1-Score<0.95, not detected: ESBL gene was not included in the bins produced by the tool, chromosome: chromosome-derived ESBL gene was included in the bins generated by the tool. (B) F1-score value distribution for all bins containing plasmid-derived ESBL genes. (C) Precision vs Recall plot for all bins containing a plasmid-derived ESBL gene.
